# Repulsive electrostatic interactions modulate dense and dilute phase properties of biomolecular condensates

**DOI:** 10.1101/2020.10.29.357863

**Authors:** Michael D. Crabtree, Jack Holland, Purnima Kompella, Leon Babl, Noah Turner, Andrew J. Baldwin, Timothy J. Nott

## Abstract

Liquid-like membraneless organelles form via multiple, weak interactions between biomolecules. The resulting condensed states constitute novel solvent environments inside eukaryotic cells that partition biomolecules and may favour particular biochemical reactions. Here we demonstrate that, in addition to attractive interactions, repulsive electrostatic interactions modulate condensate properties. We find that net charge modulates the formation, morphology and solvent properties of model Ddx4 condensates in cells and in vitro and that a net negative charge is conserved across germ cell-specific Ddx4 orthologues. This conserved net charge provides a sensitivity to multivalent cations that is not observed in somatic paralogues. The disfavouring effect of a net negative charge in Ddx4 orthologues appears to be offset by increased charge patterning, indicating that fine tuning of both attractive and repulsive interactions can create responsive solvent environments inside biomolecular condensates.

---

Condensation of biomolecules into phase separated bodies provides a reversible way for cells to compartmentalise various processes. Some of these biomolecular condensates are constitutive in eukaryotic cells [1], whereas others only occur in specific cell types or in response to certain environmental stimuli [2–8]. While a condensate may contain hundreds of different components [4], its composition is typically dominated by a small number of major constituents that define the condensate identity and likely control its overall biochemical and biophysical properties [1,2,5,9]. Relating interactions of high abundance condensate constituents to macroscopic condensate properties poses a significant challenge [10–13], but would greatly enhance both our understanding of how cells utilise intracellular phase separation and our ability to engineer orthogonal membraneless compartments.

Molecular interactions can occur over short- or long-ranges and define the stability of a molecule in solution. Short-range interactions manifest over a distance of a few angstroms and can be within a molecule – for example attractive interactions can overcome the entropic loss of constraining a protein’s conformational freedom, allowing it to fold – or they can be between molecules – for example linking the defined binding sites of two folded proteins. Long-range interactions occur over a distance of a few to tens of angstroms and can steer molecules and their binding sites towards or away from each other. For example, attraction of oppositely charged surfaces can enhance association rates [14], while repulsion of like-charged surfaces is known to influence the aggregation of proteins [15,16], and the longterm stability of colloidal dispersions like milk and paints [17,18].

Unlike bimolecular interactions between two folded proteins, where the short-range interacting residues are clustered into defined interaction surfaces, liquid-liquid phase separation relies on short-range interactions between multiple molecules [19]. Consequently, the attractive interaction sites need to be appropriately spaced to allow the formation of an intermolecular network, and have local dissociation rates that permit molecule movement within the network [20]. Intrinsically disordered regions of proteins (IDRs) provide a flexible backbone to promote network formation and phase separation through spacing interactions along the chain [21].

Compared to folded proteins, IDRs are enriched in charged residues [22]. Due to the conformational flexibility of the disordered chain, the relative 3-dimensional location of these charges is highly dynamic, allowing charge-charge interactions to occur between various, fluctuating regions of the IDR. Patterning charges into segments of high local net charge can promote interactions between oppositely charged regions [23] and, consequently, promote phase separation [24]. In cases where a difference in the number of positive and negative charges is present (i.e. when the protein has a net charge), there will be a subset of residues that are unable to interact with one of opposite charge and are instead exposed to the solvent, adding frustration into the energy landscape of the chain [25,26]. While the relative 3-dimensional stability of colloids and folded proteins provides the structure for homogenously distributed or clearly defined patches of surface charge, the frustrated excess charge in IDRs is dynamic. For a bimolecular complex with a single, defined binding interface, converting a defined region of charge on one of the molecules to a dynamic one can reduce the impact of long-range electrostatic attraction [27]. However, phase separation relies on multiple chains with multiple interaction sites, making the impact of a dynamic, effective surface charge less clear.

Here, using Ddx4 protein as a model system, we examine the effects of net charge and long-range electrostatic interactions on the formation and properties of biomolecular condensates (Fig. 1A). We find that nearly all aspects of condensate behaviour are modulated by the net charge of the primary condensate constituent.

**Figure 1.**
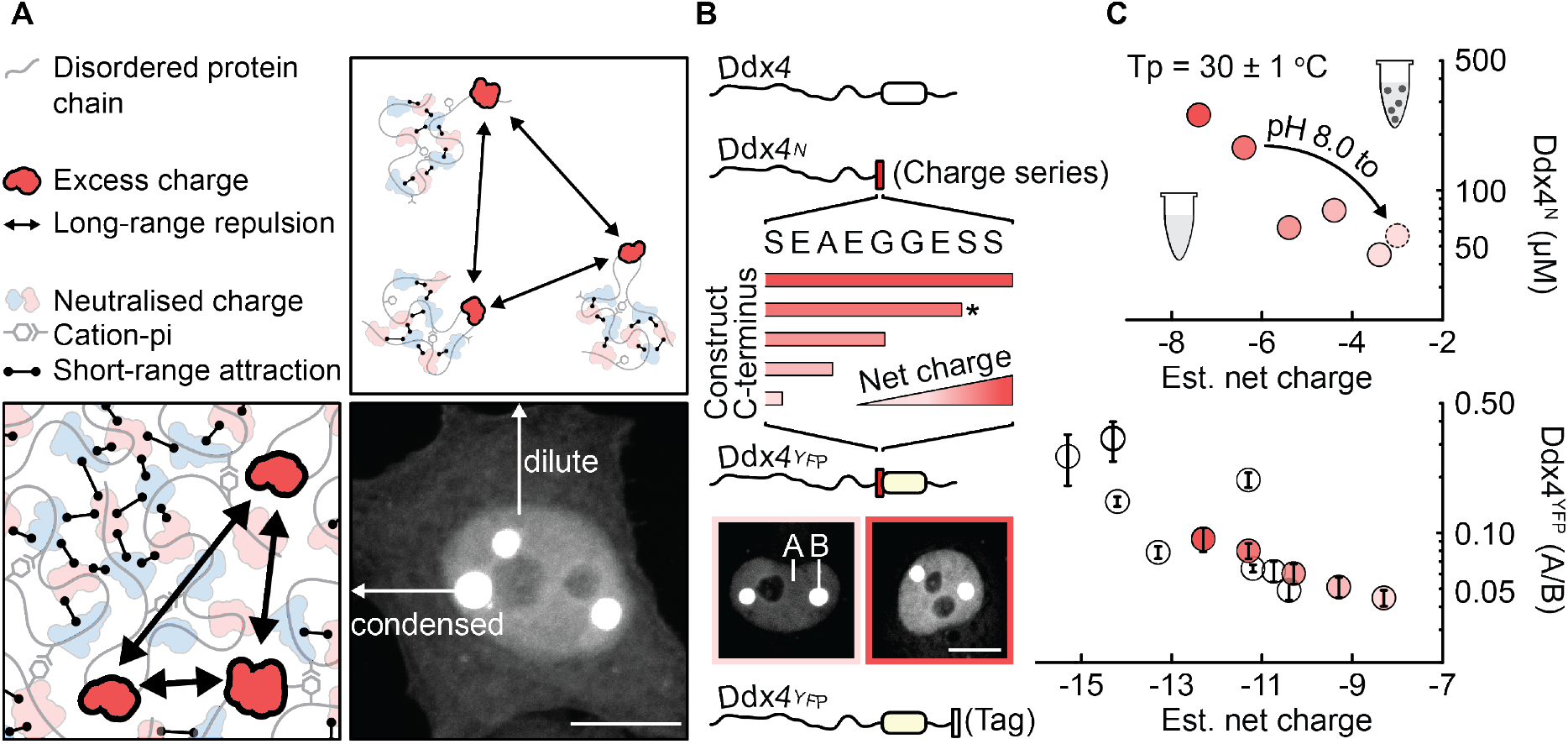
Net charge modulates long-range electrostatic repulsion and Ddx4 phase separation propensity. **(A)** The structural flexibility of intrinsically disordered regions allows residues of opposite charge to rearrange and interact, lowering the overall free energy of the chain. However, the presence of a net charge results in excess charges that cannot be satisfied through an interaction with an oppositely charged amino acid sidechain. For negatively charged Ddx4, association of dilute protein chains into a dense condensate is therefore dictated by a balance of long-range electrostatic repulsions – between excess like-charges – and shortrange attractions (e.g. between arginine and phenylalanine residues or oppositely charged residues). Scale bar 10 μm. **(B)** Slight alterations in the length of the N-terminal disordered region of Ddx4 (Ddx4^N^) reduce the net charge of the construct (top), creating a series of constructs with different estimated net charges. When expressed in HeLa cells using a construct where the DEAD-box helicase domain was replaced with YFP, the same sequence alterations led to a difference in dilute (A) and dense (B) phase fluorescence intensities, as indicated in the micrographs. Scale bar 10 μm. Further modulation of Ddx4^YFP^ net charge was achieved through the addition of short peptide tags to the C-terminus of Ddx4^YFP^(−11.3), indicated with the asterisk. See Fig. S1C for peptide tag amino acid sequences. **(C)** A lower concentration of protein was required to observe phase separation at 30 ± 1°C for Ddx4^N^ charge series constructs with lower estimated net charges (top). This was also observed when the length of the construct was kept constant (amino acids 1-236, indicated with asterisk) and pH was used to adjust the estimated net charge. The same destabilising effect of a high net charge on droplet stability was observed for Ddx4^YFP^ constructs expressed in HeLa cells, as indicated by a reduction in the difference in fluorescence intensity between the nucleoplasm (dilute phase) and condensates (dense phase) (A/B) (bottom). This was observed both when modulating net charge through the charge series sequence alterations, or through adding short peptide tags at the C-terminus of the Ddx4^YFP^(−11.3) construct. Error bars represent the standard deviation.

Ddx4 is an essential protein component of membraneless organelles termed nuage in mammals, P-granules in nematodes, and nuage, pole plasm and polar granules in flies [28–34]. Structurally, human Ddx4 and its orthologous sequences comprise two intrinsically disordered regions that flank a central DEAD-box RNA helicase domain. Assuming that the net charge of a protein at physiological pH is dominated by the number of its positively charged (arginine, R, and lysine, K) and negatively charged (aspartate, D, and glutamate, E) amino acids, we investigated the simple net charge (RK-DE) of Ddx4 orthologues annotated on the Ensembl Genome Browser [35] (Fig. S1A). These sequences, predominantly from mammals and fish, showed a net negative charge centred around −10. A smaller set of Ddx4 orthologues, chosen to give a broader view of the phylogenetic tree, showed a similar trend with a net negative charge centred around −11.

The conservation of a net negative charge in Ddx4 orthologues suggested that it may be functionally relevant. We therefore decided to investigate the impact of net negative charge on the phase separation propensity of the N-terminal disordered region of human Ddx4, Ddx4^N^. This region is sufficient to drive phase separation of Ddx4 [24] and provided an opportunity to modulate the net charge through making minimal adjustments to the length of the chain. Five constructs were produced with predicted net charges at pH 8 spanning from – 3.4 (amino acids 1-229) to −7.4 (amino acids 1-238) (Fig. 1B). Importantly, these constructs resulted in a significant titration in net charge, as confirmed with electrophoretic mobility measurements (Fig. S1B), with minimal incremental impact on the total construct length, fraction of charged residues, valency (number and distribution of arginine and phenylalanine residues) and charge patterning. Using differential interference contrast (DIC) microscopy, we monitored the concentration of each construct that was required to phase separate at 30 ± 1°C as a proxy for condensate stability. As the net charge of the protein was reduced, the concentration required to phase separate decreased, indicating that a more neutral net charge promotes phase separation of Ddx4^N^ (Fig. 1C). To confirm that our strategy for titrating Ddx4 net charge was not altering the phase separation propensity by removing interactions at the C-terminal region of the protein, we kept the length of the construct constant (236) and adjusted the pH to 6.5. This resulted in a reduction in protein concentration that was similar to that observed with the comparably charged 1-229 construct.

To determine whether the net charge of Ddx4 would also impact its phase separation in cells, we utilised a fluorescently labelled Ddx4 mimic, in which the DEAD-box helicase domain was replaced by mCitrine (Ddx4^YFP^) [24]. Net charge modulation of Ddx4^YFP^ was performed in the same manner as Ddx4^N^, creating five constructs with an estimated net charge of −8.3 to −12.3 at pH 7.4 (Fig. 1B). Importantly, while the constructs maintained the same change in net charge, the addition of YFP and the C-terminal disordered region of Ddx4 resulted in the net charge alterations occurring in the middle of protein sequence rather than at one end. Consistent with the Ddx4^N^ data in vitro, a decrease in the intensity in the dilute phase relative to the dense phase was observed upon reducing the net charge of Ddx4^YFP^, indicating that a more neutral net charge stabilises Ddx4 condensates in cells (Fig. 1C, Fig. S1D).

Next, we took advantage of the relatively highly charged nature of several commonly used short peptide tags and inserted them at the C-terminus of Ddx4^YFP^(−11.3) (Fig. 1B and S1C). This allowed us to further modulate the net charge of Ddx4^YFP^ via an orthogonal mutational approach and without making any adjustments to the N-terminal disordered region of the protein that is known to drive phase separation. Together, these results show that formation and stability of Ddx4 condensates are modulated by a balance of long-range electrostatic repulsion, mediated by the overall chain net charge, and favourable short-range interactions, predominantly between arginine and phenylalanine side chains [24,36]. The electrostatic nature of both the long-range repulsive and short-range attractive interactions means that both will be affected by the screening activity of ions in the environment. Screening of long-range electrostatic repulsion will favour phase separation while screening of short-range attractive interactions will disfavour phase separation. The impact of ions in the environment therefore depends on the how they adjust the balance of these two opposing forces (Fig. 2A).

**Figure 2.**
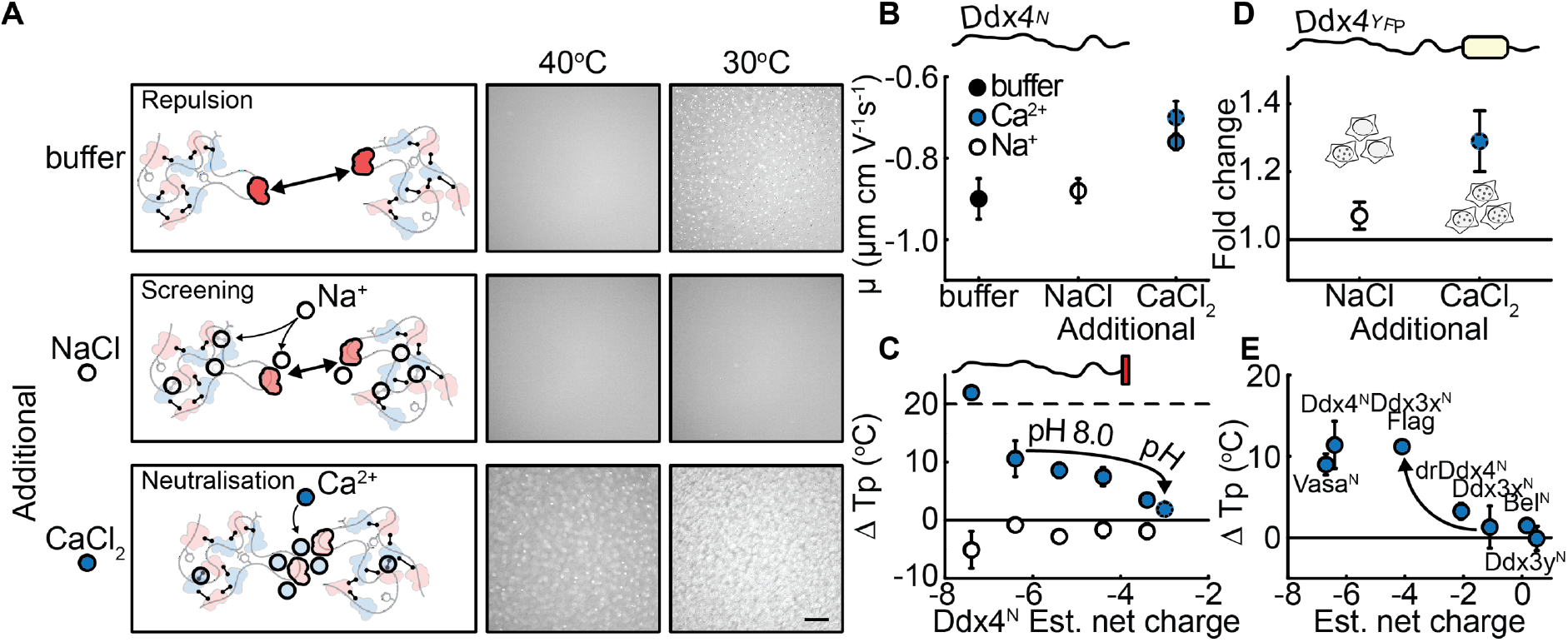
Multivalent salts alter the balance between screening short- and long-range electrostatic interactions. **(A)** Addition of NaCl screened the short-range attractive interactions of Ddx4 more than long-range repulsions. The balance was shifted for the divalent ion calcium, with long-range repulsion impacted more than short-range attraction. We suggest that this difference is due to correlations between the multivalent calcium ions that result in an increased amount of calcium surrounding the excess negative charges in Ddx4. As the calcium concentration is increased, this neutralises the net charge of Ddx4. Schematic symbol key as in Fig 1A Micrographs indicate the impact of an additional 10 mM ionic strength (I) of each salt on condensate stability. Scale bar 100 μm **(B)** Electrophoretic mobility (μ) measurements indicated that CaCl_2_ neutralises the effective surface charge of Ddx4^N^ in a titratable manner, whereas an equivalent concentration or ionic strength (I) of NaCl does not (10 mM NaCl = 10 mM I; 3.3 mM CaCl_2_ = 10 mM I; 10 mM CaCl_2_ = 30 mM I (dashed outline)). Errors represent the standard error of the mean (N = 3-4). **(C)** Addition of CaCl_2_ increased the transition temperature of Ddx4^N^, with greater differences observed for constructs with larger net charges. In contrast, addition of an equivalent ionic strength of NaCl consistently reduced the transition point temperature (Tp) of Ddx4^N^ constructs, suggesting that the charge neutralisation observed with CaCl_2_ was responsible for stabilising the condensates. The dashed line indicates the maximum difference measurable in our assay and the solid line at 0 represents the Tp calculated in buffer for each construct. Errors represent the standard deviation. **(D)** The number of HeLa cells with Ddx4^YFP^ condensates increased after exposure of the cells to extracellular CaCl_2_ (10 mM CaCl_2_ = 30 mM I) compared to an equivalent ionic strength of NaCl (30 mM NaCl = 30 mM I). Error bars represent the standard error of the mean (N=3 with >50,000 cells analysed per repeat). **(E)** Differences in transition temperature after CaCl_2_ addition (3.3 mM CaCl_2_ = 10 mM I) were dependent on the net charge of Ddx4^N^ orthologues and the paralogous Ddx3^N^ orthologues. The solid line at 0 represents the Tp calculated in buffer + 10 mM NaCl (10 mM I) for each construct. Error bars represent the standard deviation.

We have previously shown that addition of NaCl to Ddx4^N^ inhibits phase separation [24], indicating that NaCl has a greater impact on screening the attractive cation-pi interactions, than it does on reducing long-range electrostatic repulsion. This trend was also borne out by adding another monovalent salt, KCl (Fig. S2A). While the impact of monovalent salts can be described by Debye-Huckel like behaviour, salts containing multivalent ions are known to behave differently [37]. For example, multivalent ions can promote the phase separation and aggregation of polyelectrolytes, such as DNA [38], via a mechanism known as charge inversion [39]. Essentially, the addition of multivalent ions titrates the charge of the polyelectrolyte, eventually leading to a change in sign [40]. To determine if Ddx4 had a differential sensitivity to mono- and divalent ions, we first returned to measuring the electrophoretic mobility of Ddx4^N^(−6.4) in the presence of additional, biologically relevant ions (Fig. 2B). Low concentrations of additional NaCl (10 mM) did not alter the measured electrophoretic mobility. In contrast, addition of ionic strength (3.3 mM) or concentration (10 mM) matched amounts of CaCl_2_ resulted in a titratable shift in electrophoretic mobility towards zero, suggesting that CaCl_2_ was indeed reducing the effective surface charge of Ddx4^N^. We next asked whether this change in effective surface charge impacted the phase separation of Ddx4^N^ by measuring the transition temperature of a fixed protein concentration in different salt conditions (Fig. 2C). Consistent with our previous data, addition of NaCl resulted in a reduction in transition temperature for all Ddx4^N^ constructs. CaCl_2_, on the other hand, increased the transition temperature of all constructs, with larger effects observed as the net charge of the construct became more negative. Remarkably, this phase separation promoting effect of CaCl_2_ was also observed in HeLa cells (Fig. 2D, Fig. S2B).

To see if the sensitivity to calcium was a unique property of Ddx4, we investigated the impact of CaCl_2_ on the transition temperature of both Ddx4 orthologues and their somatic Ddx3 paralogues (Fig. 2E). Addition of CaCl_2_ to the N-terminal disordered region of drosophila Ddx4 (Vasa^N^) elicited an increase in transition temperature that was similar to human Ddx4^N^, while a smaller difference was observed for zebrafish Ddx4 (drDdx4^N^). Ddx3 orthologues were all minimally impacted by addition of CaCl_2_, suggesting a difference in sensitivity between the two paralogues. However, we had previously noticed a decrease in CaCl_2_ sensitivity when the net charge of Ddx4 was reduced (Fig. 2C), suggesting that excess charge is required to respond to calcium. Unlike Ddx4, which has a conserved net negative charge, Ddx3 orthologues had net charge that was closer to neutral, which we postulated may be the cause of the reduced CaCl_2_ sensitivity. We therefore modified the estimated net charge of human Ddx3x by cloning a Flag tag at the C-terminus of the Ddx3x N-terminal disordered region (Ddx3x^N^-Flag). This modified the estimated net charge from – 1.1 to −4.1 and made the phase separation of the protein sensitive to calcium, altering the transition temperature in the same manner as was observed for Ddx4 (Fig. 3E). Engineering a sensitivity to calcium in Ddx3x by modulating the net charge indicated that charge inversion via multivalent ions could be a general feature of charged, phase separating proteins.

**Figure 3.**
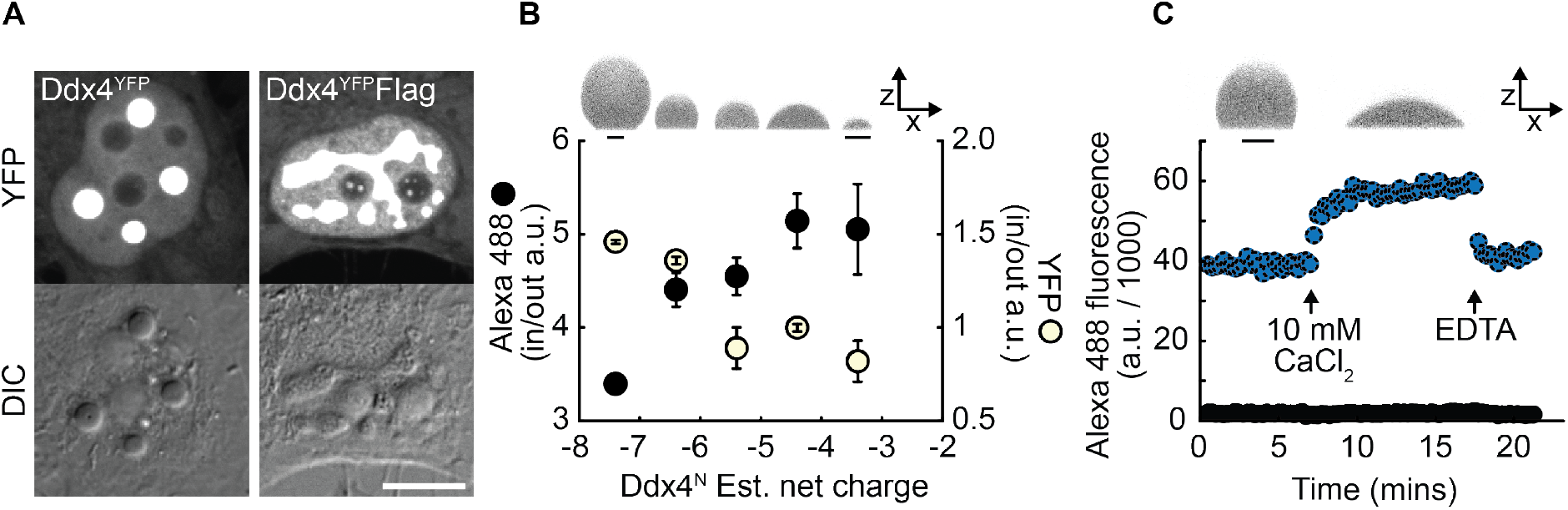
Net charge modulates the morphology and solvent properties of Ddx4 droplets. **(A)** Addition of a Flag tag (DYKDDDDK) at the C-terminal end of Ddx4^YFP^(−11.3) increased the net negative charge by −3 and led to changes in condensate morphology and fluorescence intensity in HeLa cells. Scale bar 10 μm. **(B)** In vitro xz projections of Ddx4^N^ charge series droplets sitting on siliconized glass coverslips (top). Alexa 488 was included to increase contrast. Scale bars indicate 5 μm. Common scale bar for Ddx4^N^ −7.4 to −3.4. Differences in fluorescence intensity ratio between the inside and outside of Ddx4 condensates for alexa 488 (black circles) and YFP (yellow circles) suggests that net charge alters the solvent properties of the dense phase. Error bars represent the standard deviation. **(C)** Addition of CaCl_2_ promotes the wetting of Ddx4^N^ droplets on siliconised glass coverslips and increases the maximum fluorescence intensity of alexa 488, indicating that CaCl_2_ is modulating the droplet properties in a similar manner to that observed with net charge. Scale bar indicates 5 μm (common to both droplets).

While investigating the impact of net charge on Ddx4^YFP^ phase separation in HeLa cells, we noticed that the morphology of condensates formed from Ddx4^YFP^ tagged with Flag and Myc sequences were unusual (Fig. 3A and Fig. S3A). Rather than the spherical droplets observed for Ddx4^YFP^ without a peptide tag, the fluorescence intensity of these condensates was reduced and they appeared to wet/spread across the nucleoplasm (i.e. relative to its volume, the amount of droplet surface in contact with the dilute phase had increased). In contrast, addition of the slightly basic Spot-tag did not lead to significant morphological changes, indicating that the influence of the acidic tags was due to their impact on net charge, rather than through a disruption of interactions at the C-terminus of the construct. To further investigate the impact of net charge on Ddx4 condensates, the morphology of Ddx4^N^ droplets sitting on siliconized glass coverslips was visualised using confocal fluorescence microscopy and the fluorescent dye, Alexa 488, as a probe. Similar to the in cell data, increasing the net charge of Ddx4^N^ caused the condensates to become more beaded, with more of the droplet surface in contact with the dilute phase (Fig. 3B). Additionally, Alexa 488 intensity varied for each construct, with the intensity inside of Ddx4^N^ condensates reducing relative to the outside as the net charge increased. In contrast, when the protein fluorophore, mCitrine (YFP) was used as a fluorescent probe, YFP intensity inside of Ddx4^N^ condensates was reduced relative to the outside as the Ddx4^N^ net charge was decreased (Fig. 3B). The change in fluorescence intensity ratio can be explained by a change in partitioning of the fluorophore and/or by a change in the fluorescent properties of the fluorophore. As both are dependent on the solvent properties of the dense phase relative to the dilute phase – assuming the dilute phase solvent properties were similar for all constructs – the change in ratio with net charge indicated a change in solvent properties in the Ddx4^N^ condensates. Given that CaCl_2_ impacted the electrophoretic mobility and phase separation propensity in a similar manner to reducing the construct net charge, we decided to investigate the salt’s impact on Ddx4^N^ condensate morphology and solvent properties. Changes in morphology and alexa488 intensity were observed in equilibrium and time-resolved experiments, with CaCl_2_ causing both the droplet wetting (Fig. 3C, Fig. S3B and C) and dense phase alexa488 fluorescence to increase (Fig. 3C, Fig. S3D) in a comparable manner to altering Ddx4^N^ charge via mutation. As was observed with the transition temperature experiments (Fig. 2E), this sensitivity to calcium was not present in Ddx3x^N^, but was displayed after engineering its net charge through the addition of a Flag tag (Fig. S3C).

Ddx4 orthologues have a conserved net negative charge that is not present in somatic Ddx3 paralogues, indicating that it may be functionally relevant (Fig. S4). This conservation of charge reduced the phase separation propensity of Ddx4 and impacted the morphology, solvent properties and salt sensitivity of Ddx4 condensates. From a functional perspective, Ddx4 is a major constituent of the phase separated compartment, nuage. Without Ddx4, nuage does not form [34], indicating that the ability of Ddx4 to phase separate is crucial to the condensate’s formation. It is therefore surprising that Ddx4 conserves a sequence feature that limits its ability to phase separate, and suggests that the conservation of net charge may be for its effects on the solvent, morphological and/or salt sensitivity properties of Ddx4 condensates. Nuage abundance, location and morphology changes throughout spermatogenesis [41], suggesting that it is under tight regulation and that modulation of nuage is required for appropriate germ cell maturation. Interestingly, nutritional deficiency of several multivalent ions impacts male fertility [42] and levels of calcium fluctuate during spermatogenesis [43], however, it is not currently clear that this is linked to Ddx4 regulation.

Using a model system based on Ddx4 protein, we found that aspects of condensate behaviour, including the concentration required for phase separation, droplet stability, wetting and morphology, solvent properties, and the response to other ions in the environment, are all dependent on overall chain net charge (Fig. 4A). We suggest that a balance of both attractive and repulsive interactions confers the ability to modulate biological condensate properties (Fig. S4). Addition of net charge to the molecular grammar of phase separation could help to explain effects seen after post-translational modification of phase separating proteins [44–47], and could aid in designing synthetic phase separating systems with unique properties.

**Figure 4.**
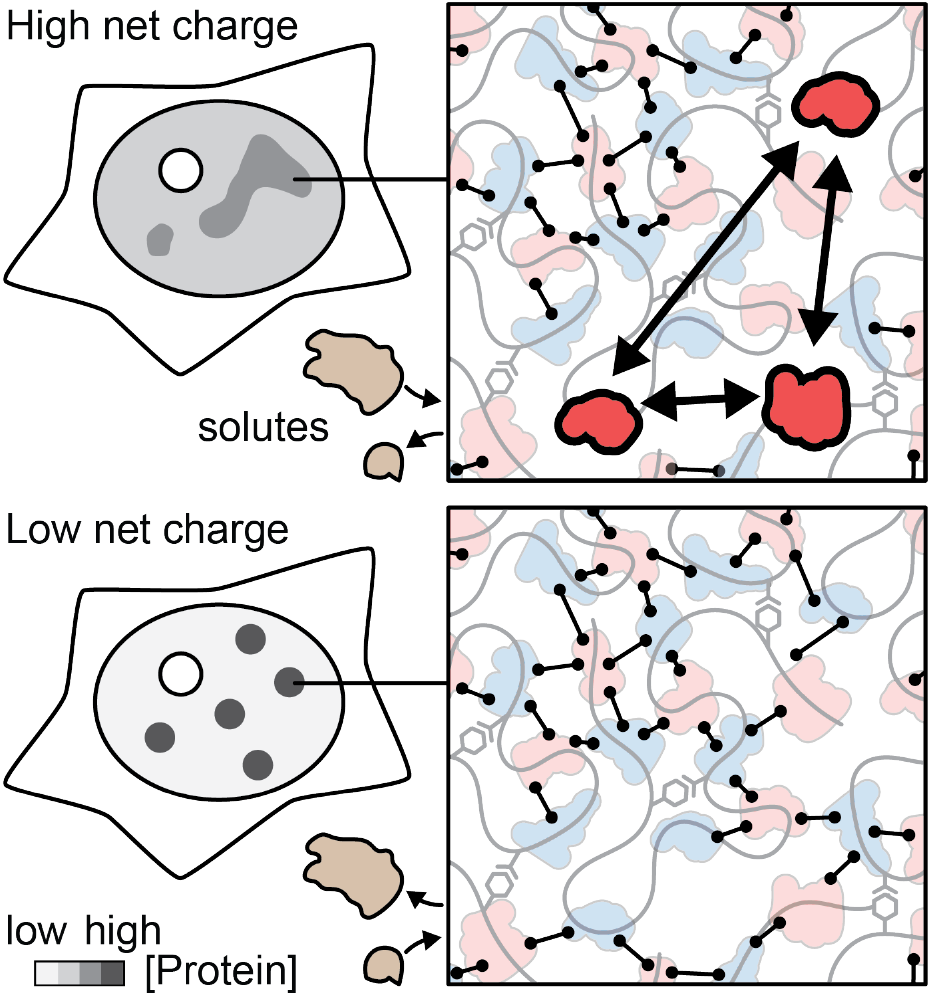
A balance of attractive and repulsive interactions modulates droplet stability, morphology and solvent properties. Schematic indicating the effects of net charge on a phase separating protein expressed in cells. Relative to a more neutral protein (bottom), proteins with high net charge (top) required a greater concentration to phase separate (higher dilute phase concentration), had a lower difference in intensity between the dense and dilute phases, and a less spherical morphology in cells. The ratio of solute intensities in the dense and dilute phases was also altered, indicating the influence of net charge on the solvent properties of the dense phase. Schematic symbol key as in Fig 1A.

## Acknowledgements

Funding for this work came from a Sir Henry Dale Fellowship jointly funded by the Wellcome Trust and the Royal Society (Grant Number 202320/Z/16/Z) awarded to T.J.N. M.D.C. thanks New College, Oxford, for additional support. M.D.C. and T.J.N. thank D.G.A.L. Aarts for discussion and the use of instrumentation, Micron Oxford for microscopy support and M. Maj for support with flow cytometry experiments. T.J.N. thanks F. Barr and members of the Barr group for the gift of HeLa cells and support with tissue culture.

## Materials and Methods

### Protein expression and purification

DNA sequences for Ddx4 and Bel constructs were generated by PCR and subcloned into pET SUMO vectors. Transformed BL-21 codon plus RIL *E. coli* cells were grown at 37°C to an optical density of 0.6-1 in terrific broth and induced with 0.5 mM IPTG. Protein expression was left to occur overnight at 22°C, 180 rpm. To reduce protein degradation during sonication and affinity purification, cell pellets were resuspended in 20 mM sodium phosphate, 10 mM imidazole, 6 M guanidinium hydrochloride (GdmCl), pH 7.4. Sonicate supernatants were loaded onto Ni-NTA agarose resin and incubated for >1 hour at 4°C. After washing with 20 mM sodium phosphate, 10 mM imidazole, pH 7.4, the bound SUMO-tag was removed by the protease ULP-1, and cleaved Ddx4 was eluted from the resin supernatant. The protein was further purified by size exclusion chromatography, using an elution buffer of 20 mM Tris, 300 mM NaCl, 5 mM TCEP, pH 8 at 22°C.

GST tagged protein constructs (Ddx3x, Ddx3y, Vasa, YFP) were generated by subcloning sequences of interest into a modified pETM-30 vector containing the pGEX-2T-TEV site and pProEx multiple cloning site. Transformed BL-21 codon plus RIL *E. coli* cells were grown at 37°C to an optical density of 0.6-1 in terrific broth and induced with 0.5 mM IPTG. Protein expression was left to occur for 4 hours at 37°C or overnight at 22°C. Cell pellets were typically resuspended in buffer containing 20 mM Tris pH 8 at 22-25°C, 300 mM NaCl, 2 mM DTT and EDTA-free protease inhibitor tablets (Roche). Sonicate supernatants were loaded onto GST-4B resin (Amersham) and incubated for >1 hour at 4°C. Resin was typically washed with 20 mM Tris pH 8, 1.5 M NaCl, 2 mM DTT and then exchanged into 20 mM Tris pH 8 at 22, 300 mM NaCl, 2 mM DTT, 2 mM EDTA. The GST-tag was removed by TEV protease, and cleaved protein was eluted from the resin supernatant. The protein was further purified by size exclusion chromatography, using an elution buffer of 20 mM Tris, 300 mM NaCl, 1 mM TCEP, pH 8 at 22-25°C.

Protein purity and identities were confirmed by SDS-PAGE and mass spectrometry, respectively.

### HeLa Cell culture and transfection

HeLa cells were cultured as previously described [9,24]. Briefly, HeLa cells were grown on 22 mm diameter #1.5 glass coverslips (Agar Scientific) in growth media (high glucose DMEM (Gibco™) containing 10% FBS (Sigma)) at 37°C and 5% CO_2_. Protein constructs were expressed in HeLa cells from pcDNA 3.1+ (Invitrogen) plasmids by transient transfection using the TransIT^®^-LT1 Transfection Reagent (Mirus). Transfections used 0.5 – 1 μg plasmid DNA per coverslip and were carried out according to the manufacturer’s instructions. Transfected cells were fixed approximately 24 hours after transfection.

### Protein Sequences

Amino acid sequences for the constructs used in this work are shown below. Sequences correspond to the longest length constructs, with mutated regions shown in red and remnants from cloning in blue. Underlined residues indicate fluorophore sequences. Highlighted regions indicate the location of peptide tags.

Ddx4^N^ (Uniprot: Q9NQI0; 1-238)

GAMGSMGDEDWEAEINPHMSSYVPIFEKDRYSGENGDNFNRTPASSSEMDDGPSRRDHFMKSGFASGRNFGNRDAGECNKRDNTSTMGGFGVGKSFGNRGFSNSRFEDGDSSGFWR ESSNDCEDNPTRNRGFSKRGGYRDGNNSEASGPYRRGGRGSFRGCRGGFGLGSPNNDL DPDECMQRTGGLFGSRRPVLSGTGNGDTSQSRSGSGSERGGYKGLNEEVITGSGKNSW KSEAEGGESSD

Ddx3x^N^ (Uniprot: Q9NQI0; 1-137)

GAMGSMSHVAVENALGLDQQFAGLDLNSSDNQSGGSTASKGRYIPPHLRNREATKGFYDKDSSGWSSSKDKDAYSSFGSRSDSRGKSSFFSDRGSGSRGRFDDRGRSDYDGIGSRGD RSGFGKFERGGNSRWCDKSDEDDWS[FLAG]

Ddx3y^N^ 135 (Uniprot: O15523; 1-135)

GAMGSMSHVVVKNDPELDQQLANLDLNSEKQSGGASTASKGRYIPPHLRNREASKGFHDKDSSGWSCSKDKDAYSSFGSRDSRGKPGYFSERGSGSRGRFDDRGRSDYDGIGNRERP GFGRFERSGHSRWCDKSVEDDWS

Vasa^N^ (Uniprot: P09052; 1-192)

GAMGSMSDDWDDEPIVDTRGARGGDWSDDEDTAKSFSGEAEGDGVGGSGGEGGGYQGGNRDVFGRIGGGRGGGAGGYRGGNRDGGGFHGGRREGERDFRGGEGGFRGGQGGSR GGQGGSRGGQGGFRGGEGGFRGRLYENEDGDERRGRLDREERGGERRGRLDREERGGERGERGDGGFARRRRNEDDINNNNNIV

Bel^N^ (Uniprot:Q9VHP0; 1-252)

GAMGSMSNAINQNGTGLEQQVAGLDLNGGSADYSGPITSKTSTNSVTGGVYVPPHLRGGGGNNNAADAESQGQGQGQGQGFDSRSGNPRQETRDPQQSRGGGGEYRRGGGGGGR GFNRQSGDYGYGSGGGGRRGGGGRFEDNYNGGEFDSRRGGDWNRSGGGGGGGRGFGRGPSYRGGGGGSGSNLNEQTAEDGQAQQQQQPRNDRWQEPERPAGFDGSEGGQSAGGNRSYNNRGERGGGGYNSRWKEGGGSNVDYT

Ddx4^YFP^

MGDEDWEAEINPHMSSYVPIFEKDRYSGENGDNFNRTPASSSEMDDGPSRRDHFMKSGFASGRNFGNRDAGECNKRDNTSTMGGFGVGKSFGNRGFSNSRFEDGDSSGFWRESSNDCEDNPTRNRGFSKRGGYRDGNNSEASGPYRRGGRGSFRGCRGGFGLGSPNNDLDPDECMQRTGGLFGSRRPVLSGTGNGDTSQSRSGSGSERGGYKGLNEEVITGSGKNSWKSEAEGGESSD<u>MVSKGEELFTGVVPILVELDGDVNGHKFSVSGEGEGDATYGKLTLKFICTTGKLPVPWPTLVTTFGYGLMCFARYPDHMKQHDFFKSAMPEGYVQERTIFFKDDGNYKTRAEVKFEGDTLVNRIELKGIDFKEDGNILGHKLEYNYNSHNVYIMADKQKNGIKVNFKIRHNIEDGSVQLADHYQQNTPIGDGPVLLPDNHYLSYQSKLSKDPNEKRDHMVLLEFVTAAGITF</u>STYIPGFS GSTRGNVFASVDTRKGKSTLNTAGFSSSQAPNPVDDESWD(GSGSG[tag])

Ddx4^CFP^

MGDEDWEAEINPHMSSYVPIFEKDRYSGENGDNFNRTPASSSEMDDGPSRRDHFMKSGFASGRNFGNRDAGECNKRDNTSTMGGFGVGKSFGNRGFSNSRFEDGDSSGFWRESSNDCEDNPTRNRGFSKRGGYRDGNNSEASGPYRRGGRGSFRGCRGGFGLGSPNNDLDPDECMQRTGGLFGSRRPVLSGTGNGDTSQSRSGSGSERGGYKGLNEEVITGSGKNSWKSEAEGGES<u>MVSKGEELFTGVVPILVELDGDVNGHKFSVSGEGEGDATYGKLTLKFICTTGKLPVPWPTLVTTLTWGVQCFARYPDHMKQHDFFKSAMPEGYVQERTIFFKDDGNYKTRAEVKFEGDTLVNRIELKGIDFKEDGNILGHKLEYNAISDNVYITADKQKNGIKANFKIRHNIEDGSVQLADHYQQNTPIGDGPVLLPDNHYLSTQSKLSKDPNEKRDHMVLLEFVTAAGIT</u>FSTYIPG FSGS TRGNVFASVDTRKGKSTLNTAGFSSSQAPNPVDDESWD

YFP (mCitrine)

GAMGSMVSKGEELFTGVVPILVELDGDVNGHKFSVSGEGEGDATYGKLTLKFICTTGKLPVPWPTLVTTFGYGLMCFARYPDHMKQHDFFKSAMPEGYVQERTIFFKDDGNYKTRAEVKFEGDTLVNRIELKGIDFKEDGNILGHKLEYNYNSHNVYIMADKQKNGIKVNFKIRHNIEDGSVQLADHYQQNTPIGDGPVLLPDNHYLSYQSKLSKDPNEKRDHMVLLEFVTAAGIT

### Buffers and reagents

Unless otherwise stated, in vitro experiments were performed in a pH 8 buffer containing 20 mM Tris, 150 mM NaCl and 5 mM TCEP. This was typically achieved through mixing protein solutions (20 mM Tris, 300 mM NaCl, 5mM TCEP) with 20 mM Tris, 5 mM TCEP, which reduced both the ionic strength and protein concentration of the stock solution and promoted phase separation. For salt studies, 4x salt solutions were mixed with 2x buffer solutions (e.g. 40 mM CaCl_2_ mixed with 40 mM Tris, 10 mM TCEP). This gave a final 2x salt solution in 1x buffer that was mixed with the protein solution (e.g. providing final buffer conditions of 20 mM Tris, 150 mM NaCl, 10 mM CaCl_2_, 5 mM TCEP). To improve the accuracy of the dilutions, all mixing steps outlined above were performed with a 1:1 volume ratio.

For Hela cell experiments involving treatment with salts, cells were grown in DMEM (ThermoFisher; 31966021) with 9 % FBS (Sigma; F9665).

### Electrophoretic mobility

Electrophoretic mobility measurements were performed using a Zetasizer Nano (Malvern Pananalytical). Samples were prepared at 15 μM and filtered (0.22 μm) prior to taking the measurement. The mean electrophoretic mobility and associated errors were calculated from 2-4 independently prepared sample replicates, with each replicate consisting of at least 3 estimates.

### Net charge calculator

To calculate the net charge of each protein construct, the proportion of each ionisable group was calculated at each pH using the Henderson-Hasselbalch equation and the following pKa values; C-terminus = 3.6, ASP = 4, GLU = 4.5, HIS = 6.4, N-term = 7.8, CYS = 8.14, TYR = 9.6, LYS = 10.4, ARG = 12.5. This was multiplied by the number and sign of each respective amino acid/ionisable group that was present in the construct, with the net charge determined from the sum of these values.

### Transition temperature determination

Transition temperature measurements were typically performed as described previously [48]. Briefly, 0.22 mm thick siliconized glass coverslips (Hampton Research), buffers and protein solutions were preheated on the heating block of a thermomixer. The ionic strength of the protein solution was diluted with buffer (typically 20 mM Tris, 5 mM TCEP, pH 8) containing various concentrations of different salts before transferring to the coverslip. The imaging chamber was sealed with 0.12 mm imaging spacers (Sigma) and a second siliconized glass coverslip. Samples were transferred to a pre-heated Linkam PE120xy temperature-controlled imaging stage controlled with LinkSys software (Linkham) and imaged using a 10x differential interference contrast (DIC) objective on an Olympus BX43 microscope. Temperature ramps typically consisted of a 2°C min^-1^ reduction in temperature and were initiated at least 10°C above the transition temperature.

For analysis, twelve images of an isothermal sample were captured at the peak temperature of the ramp and used to calculate a baseline pixel intensity. Upon phase separation, condensate formation resulted in a change in observed pixel intensity. The transition temperature was then defined as the temperature at which the pixel intensity deviated by >10 standard deviations of the baseline intensity.

### Confocal fluorescence microscopy of in vitro samples

To increase contrast through fluorescence imaging, the fluorescent molecules, Alexa 488 or mCitrine (YFP), were included in all samples at a final concentration of approximately 0.6 μM and 2 μM, respectively. Images were captured at room temperature using a Leica TCS-SP5 confocal fluorescence microscope and a 63x oil immersion objective. Samples were illuminated with a 488 nm (Alexa 488) or 514 nm (YFP) laser, with power and gain settings adjusted to give mean fluorescence intensity values of the dense phase of approximately 50-70% of the maximum 16-bit depth (~32,000-46,000 a.u.). Images were typically captured with settings of 256 x 256 pixels at ~ 98 x 98 x 98 nm (XYZ) resolution, 1400 hz and a line average of 2.

Phase separation was initiated by mixing proteins solutions with a buffer of lower ionic strength, as described above, and were left for 10 mins to promote droplet growth. Phase separated solutions were then transferred to a 0.22 mm thick siliconized glass coverslip (Hampton Research), before sealing with 0.12 mm imaging spacers (Sigma) and a second siliconized glass coverslip. Samples were then left to equilibrate for approximately 50 minutes prior to imaging.

### In vitro partitioning

Image analyses were performed using bespoke procedures in Mathematica 12. 3D image stacks of sessile droplets resting on a solid substrate were collected using laser scanning confocal fluorescence microscopy as above. In the positive z-direction, optical sectioning moved out of the solid substrate into the aqueous solution containing droplets. The xy slices containing the aqueous solution were identified by first measuring the change in pixel standard deviation, per slice, in z. The maxima of the change in standard deviation of pixel intensity in z was taken to approximate the position of the solid-aqueous interface. All images in z positions greater than this value plus 3 were carried forwards. A gaussian blur was then applied, using the “GaussianFilter” function with a kernel of radius 3, to the 3D image stack composed of the aqueous phase images. The blurred images were then binarized using the “Binarize” function with the default method of Otsu’s algorithm. This generated a binary mask containing the condensed protein phase, whilst excluding the dilute protein phase. Dense phase intensity values were then extracted using the “ImageMeasurements” function from the original 3D image stack containing only the aqueous phase as the first argument, “MeanIntensity” as the second argument and an eroded version of the condensed phase mask (generated using the “Erosion” function”). Equivalently, dilute phase intensity values were extracted using the same procedure, except for the application of an eroded negative of the condensed phase mask to identify only the dilute phase. In both cases, the erosion ensured no edge effects caused by the blur of dilute phase voxels with dense phase voxels.

YFP typically generated little contrast between the dilute and dense phase, making automated droplet identification difficult. Mean pixel intensities where therefore determined from manually drawn regions of interest from at least five fields of view. Before calculating the fluorescence intensity ratio, background dense and dilute phase intensities, determined from samples lacking a fluorophore, were subtracted.

### Imaging fixed HeLa cells

HeLa cells expressing fluorescent proteins were grown on 22 mm diameter glass coverslips, washed twice with phosphate buffered saline (PBS) at 37°C and fixed with 4% paraformaldehyde (PFA) in PBS (Alfa Aesar) at 37°C, for 5 minutes. Cells were then washed three times with PBS to remove excess PFA with the first wash only containing Hoechst 33342 dye at 2 μM. Cells were incubated for 5-10 minutes in the wash solution at each step. Coverslips were mounted on microscope slides (Fisher Scientific™) using Immu-Mount (Thermo Scientific™ Shandon™) mounting medium.

Cells were imaged on an Olympus FV1000 Laser Scanning Microscope based on an Olympus IX81 inverted microscope with a motorised stage, and equipped with an Olympus PLAPON60XOSC2 (NA 1.4) oil immersion objective. Hoechst was excited with a solid state 405 nm laser and YFP (mCitrine) was excited with an argon 515 nm laser. Hoechst fluorescence was collected between (420 – 435 nm) and YFP fluorescence was collected between (530 – 650 nm). DIC images were collected using illumination from the 515 nm laser.

Unless otherwise stated, imaging data were collected as stacks of 86 Z-slices (256 x 256 pixels (xy), 0.1 μm spacing (z), 2 μs pixel^-1^ scanning speed, 12 bit depth, 2 x line averaging) for each channel. In all cases the YFP stack was collected before the Hoechst channel stack was collected. Image analysis was performed using Wolfram Mathematica (Wolfram Research, Inc., Mathematica, Version 12.0, Champaign, IL (2019)) and Fiji [49]. Figures were prepared using Fiji and Omero Figure [50].

### In cell determination of saturation concentration, partitioning and fluorescence intensity of C-terminally tagged constructs

Imaging data was collected with identical laser power and detector sensitivity so that fluorescence intensities across difference samples were directly comparable. Image analyses to extract intensities for Ddx4^YFP^ condensates and the nucleoplasm (excluding nucleoli) were performed using Fiji. First, maximum intensity z-projections were made from each stack and a histogram of pixel intensity (64 bins) was generated. The peak of the histogram (logio y-axis) in the high-intensity range (i.e. close to the maximum value) was taken as a proxy for mean condensate fluorescence intensity in that stack. Nucleoplasmic fluorescence intensity was determined as the highest frequency bin following manual removal of regions corresponding to nucleoli, Ddx4^YFP^ condensates and extra-nuclear regions. Fluorescence intensity values for Ddx4^YFP^ condensates and the nucleoplasm were then averaged from multiple stacks for the same construct.

### Real-time droplet imaging

Ddx4^N^ (amino acids 1-236) was allowed to phase separate by mixing in a 1:1 volume ratio with 20 mM Tris, 5 mM TCEP, 1.2 μM Alexa 488 (final buffer composition of 20 mM Tris, 150 mM NaCl, 5 mM TCEP, 0.6 μM Alexa 488, pH 8 at 22°C), before transferring 2 μL to a siliconized glass coverslip. To prevent evaporation during the course of the experiment, 10 μL of mineral oil was added on top of the beaded Ddx4 solution, forming a seal. To rapidly collect XYZ confocal images (approximately 15s per stack), settings of 128 x 128 pixels at ~300 x 300 x 300 nm (XYZ) resolution, 1400 hz and line average of 2 were used.

Solutions containing CaCl_2_ and EDTA were generated by mixing a 4x stock with Ddx4^N^ 1-236. For example, 20 mM Tris, 5 mM TCEP, pH 8, 40 mM CaCl_2_, 1.2 μM Alexa 488 was mixed with Ddx4^N^ 1-236 giving final conditions of Ddx4N 1-236 in 20 mM Tris, 150 mM NaCl, 5 mM TCEP, pH 8, 20 mM CaCl_2_, 0.6 μM Alexa 488. Using a 10 μL pipette tip to pierce the mineral oil film, 2 μL of this solution was then added to the solution being imaged. This ensured that all components except the additive (e.g. CaCl_2_) remained at the same concentration in the sample. Upon removing the pipette tip, the mineral oil reformed the seal, preventing any water loss through evaporation.

### Flow cytometry

HeLa cells were grown in 15 cm plates to a confluency of approximately 50% and transfected with 10 μg of DNA. Expression was allowed to proceed over a 24-hour incubation period at 37°C and 5% CO_2_, after which cells had typically reached a confluency of 70-80%. Cells were then dissociated using trypsin, pelleted and resuspended in PBS. To ensure differences in transfection efficiencies and passage number were not conflated with differences between salt conditions, resuspended cells from the same plate were split into different tubes. Cells were then pelleted and resuspended 20 mM Hepes, 137 mM NaCl, 2.7 mM KCl, pH 7.4 at 37°C, with or without additional salt (e.g. 20 mM Hepes, 137 mM NaCl, 2.7 mM KCl, 10 mM CaCl_2_). Hepes was chosen at this stage to prevent phosphate precipitation by calcium. Cells were pelleted and resuspended again before incubating at 37°C and 5% CO_2_ for 10 mins. Fixation was achieved by resuspending pelleted cells in PBS with 4% paraformaldehyde and incubating at room temperature for 10 mins, before washing cells with PBS. To concentrate cells for flow cytometry, cells were pelleted and resuspended in 100-200 μL of PBS. Cell data was collected by passing cells through an Amnis Imagestream flow cytometer. Typically, data from 100,000 cells were collected from each experimental replicate.

To identify cells with Ddx4 condensates, two parameters were used. Firstly, as formation of Ddx4 condensates occurs in a protein concentration dependent manner, we identified cells with high fluorescence intensity levels, which indicated high expression of the YFP-tagged Ddx4 construct. Secondly, the difference in protein concentration between the dense and dilute phases results in Ddx4 condensates having significantly higher fluorescence intensity than the surrounding dilute phase. We reasoned that cells with high fluorescence contrast would therefore be more likely to have condensates, whereas cells with high concentrations of YFP-tagged Ddx4 but no condensates would have a more uniform fluorescence intensity and lower contrast. By plotting fluorescence intensity against fluorescence contrast, a population of cells with high intensity and high contrast could clearly be distinguished (Figure S2). Upon investigating images of these cells captured by the Imagestream cytometer, it appeared that these cells contained condensates. Furthermore, this population of cells was not present in cells expressing only YFP, indicating that these cells contained condensates composed of Ddx4. To count cells within this population, a boundary was drawn. This reduced the number of cells from the initial 100,000 to approximately 1000. Cells within this boundary were then manually inspected to remove false positives and ensure that condensate-like features could be identified. At least three experiments were performed for each salt condition.

### Sequence analysis

We obtained amino acid sequences for Ddx4 and Ddx3x orthologues from UniProt (https://doi.org/10.1093/nar/gky1049) and Ensembl (release 102 – November 2020, https://doi.org/10.1093/nar/gkz966). Simple net charge was computed as follows: amino acids Arginine and Lysine each contributed a charge of +1 and Aspartic Acid and Glutamic Acid each contributed a charge of −1.

Alignment of orthologous Ddx4 and Ddx3x sequences was performed in Jalview [51] using the Muscle algorithm with default settings. Boundaries for protein domains (N- and C-terminal IDRs and central helicase domains) were determined from alignment of all orthologous sequence sets and exemplified by the human sequences (amino acid numbers in Table 1). Overall IDR length, aromatic content and charge patterning (kappa, calculated using localCIDER [52] were determined by first removing aligned helicase domains and then concatenating N- and C-terminal IDRs (IDR^NC^). Sequences with IDR^NC^ ≤ 100 amino acids in length typically corresponded to Ensembl entries with poor sequence coverage and were not included in the analyses. Density plots were produced via Gaussian kernel density estimation in Python with the same smoothing factor used on equivalent Ddx4 and Ddx3x data.

**Table 1.**
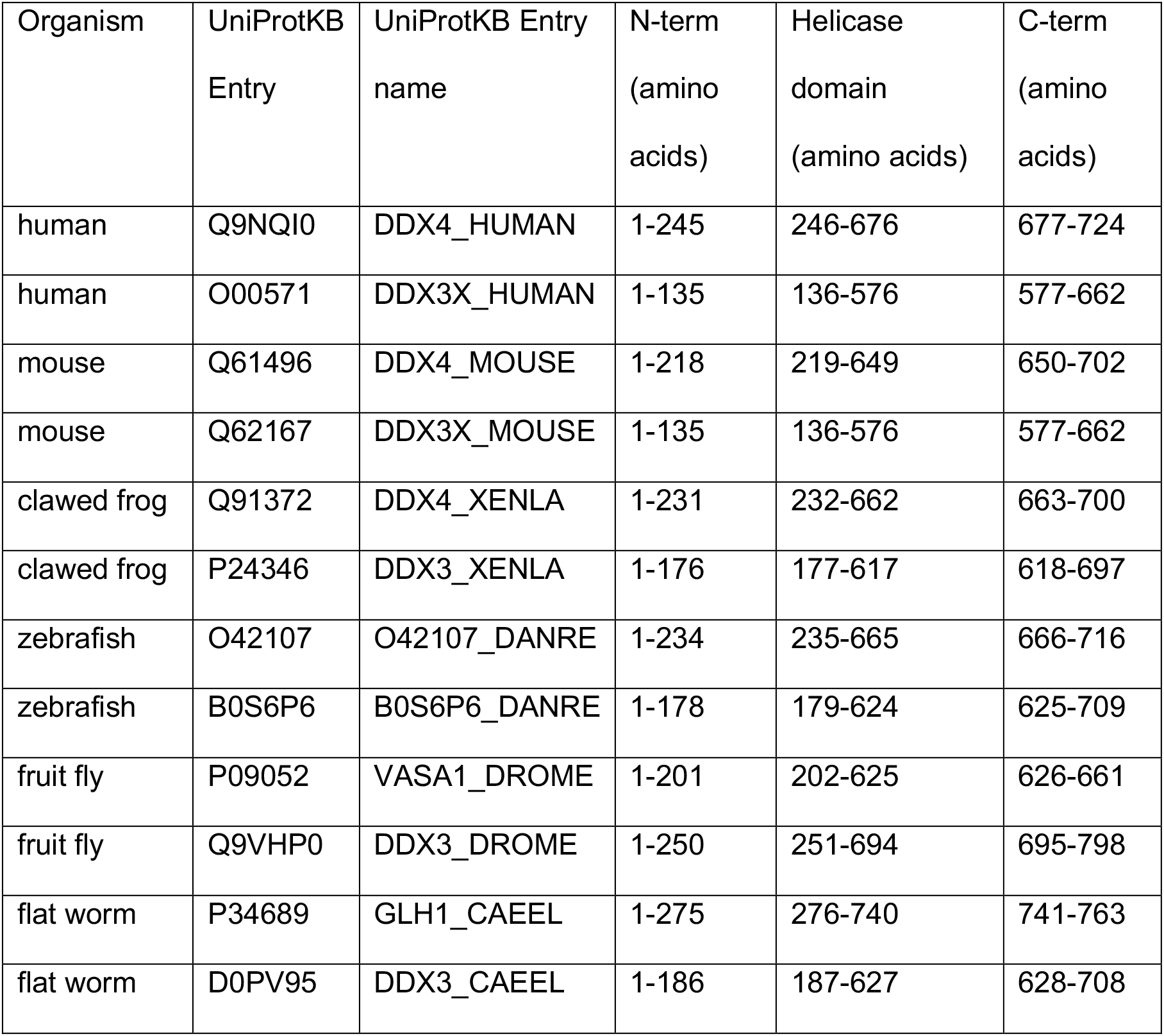
Gene names and domain boundaries used to calculate net charges displayed in Figure S4.

| Organism | UniProtKB Entry | UniProtKB Entry name | N-term (amino acids) | Helicase domain (amino acids) | C-term (amino acids) |
| --- | --- | --- | --- | --- | --- |
| human | Q9NQI0 | DDX4_HUMAN | 1-245 | 246-676 | 677-724 |
| human | O00571 | DDX3X_HUMAN | 1-135 | 136-576 | 577-662 |
| mouse | Q61496 | DDX4_MOUSE | 1-218 | 219-649 | 650-702 |
| mouse | Q62167 | DDX3X_MOUSE | 1-135 | 136-576 | 577-662 |
| clawed frog | Q91372 | DDX4_XENLA | 1-231 | 232-662 | 663-700 |
| clawed frog | P24346 | DDX3_XENLA | 1-176 | 177-617 | 618-697 |
| zebrafish | O42107 | O42107_DANRE | 1-234 | 235-665 | 666-716 |
| zebrafish | B0S6P6 | B0S6P6_DANRE | 1-178 | 179-624 | 625-709 |
| fruit fly | P09052 | VASA1_DROME | 1-201 | 202-625 | 626-661 |
| fruit fly | Q9VHP0 | DDX3_DROME | 1-250 | 251-694 | 695-798 |
| flat worm | P34689 | GLH1_CAEEL | 1-275 | 276-740 | 741-763 |
| flat worm | D0PV95 | DDX3_CAEEL | 1-186 | 187-627 | 628-708 |

**Figure S1.**
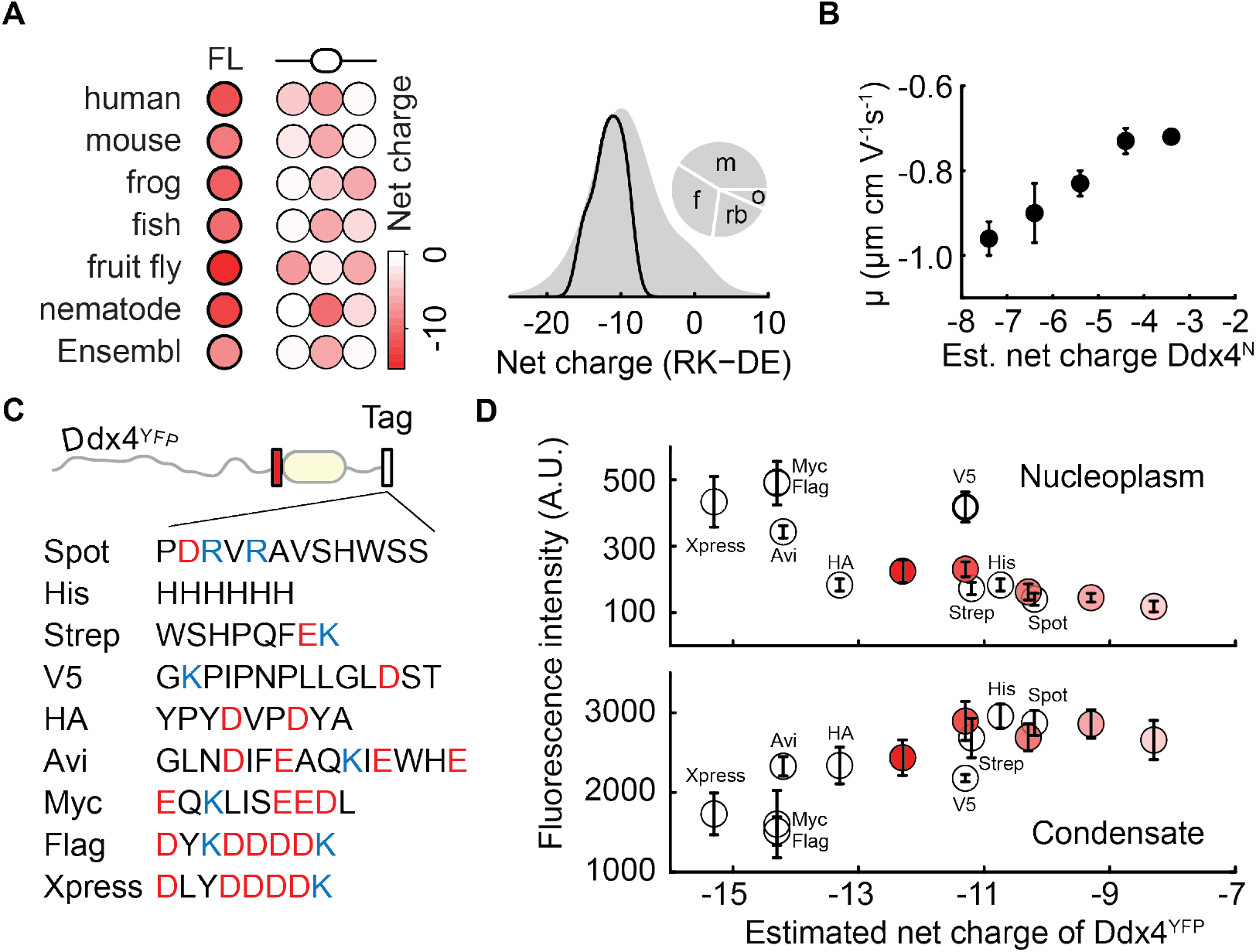
A conserved net negative charge modulates the stability of Ddx4 condensates in vitro and in cells. **(A)** The folded helicase domain (oval), and N- and C-terminal disordered regions (line) of Ddx4 all contribute either a neutral or negative charge to the full-length (FL) protein. Density distributions of Ddx4 net charge from orthologues annotated on Ensembl (grey) or a phylogenetically broad subset (black) of human, mouse, frog, fish, fruit fly and nemotode sequences. The pie chart indicates the proportion of sequences from mammals (m), bony fish (f), reptiles and birds (rb) and other species (o) in the Ensembl dataset. **(B)** Truncating the N-terminal region of Ddx4^N^ alters the observed electrophoretic mobility (μ), suggesting a change in the effective surface charge. Error bars represent the standard deviation. **(C)** Sequences of C-terminal tags that were used to modulate the net charge of Ddx4^YFP^. **(D)** Fluorescence intensity measurements for the interior and exterior of Ddx4^YFP^ condensates in HeLa cells. The ratio of these values in shown in Figure 1C. Error bars represent the standard deviation.

**Figure S2.**
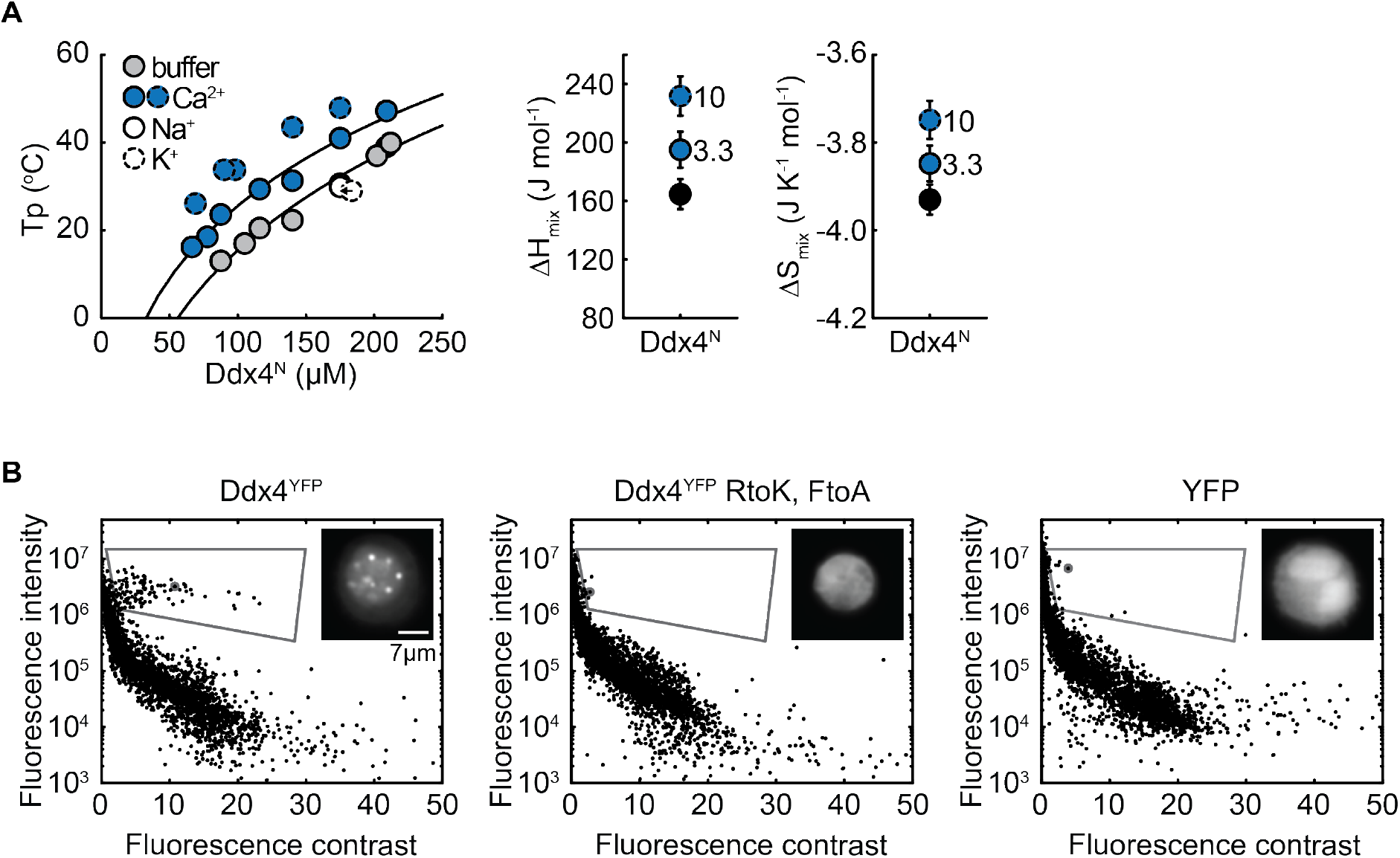
Calcium modulates the stability of Ddx4 droplets in vitro and in cells. **(A)** The temperature at which phase separation was observed for various concentrations of Ddx4^N^ was determined in 20 mM Tris, 150 mM NaCl, 5 mM TCEP (buffer) +/- additional salts (10 mM NaCl, 10 mM KCl, 3.3 mM CaCl_2_ and 10 mM CaCl_2_). To examine how the entopic and enthalpic contributions to the interaction parameter varied upon addition of CaCl_2_, data for buffer, 3.3 mM CaCl_2_ and 10 mM CaCl_2_ (markers with dashed outlines) were fit to a Flory-Huggins model of polymer phase separation, as described previously [24]. **(B)** Flow cytometry data for HeLa cells expressing Ddx4^YFP^(−11.3) indicated a subset of cells with high fluorescence intensity and contrast that were not present in cells expressing either YFP or an RtoK, FtoA mutant of Ddx4^YFP^(−11.3) that is known to reduce the phase separation propensity of Ddx4 [24]. Images taken during flow cytometry indicated that this subset was reporting on cells with condensate like features, allowing the number of cells with condensates to be determined in a high throughput manner. A grey box indicates the region where cells with condensates were identified and markers with grey outlines indicate the data point that corresponds to each cell image. Data represents a run of 10,000 cells for each construct, with approximately 1% of Ddx4^YFP^ cells falling within the region outlined with the grey box.

**Figure S3.**
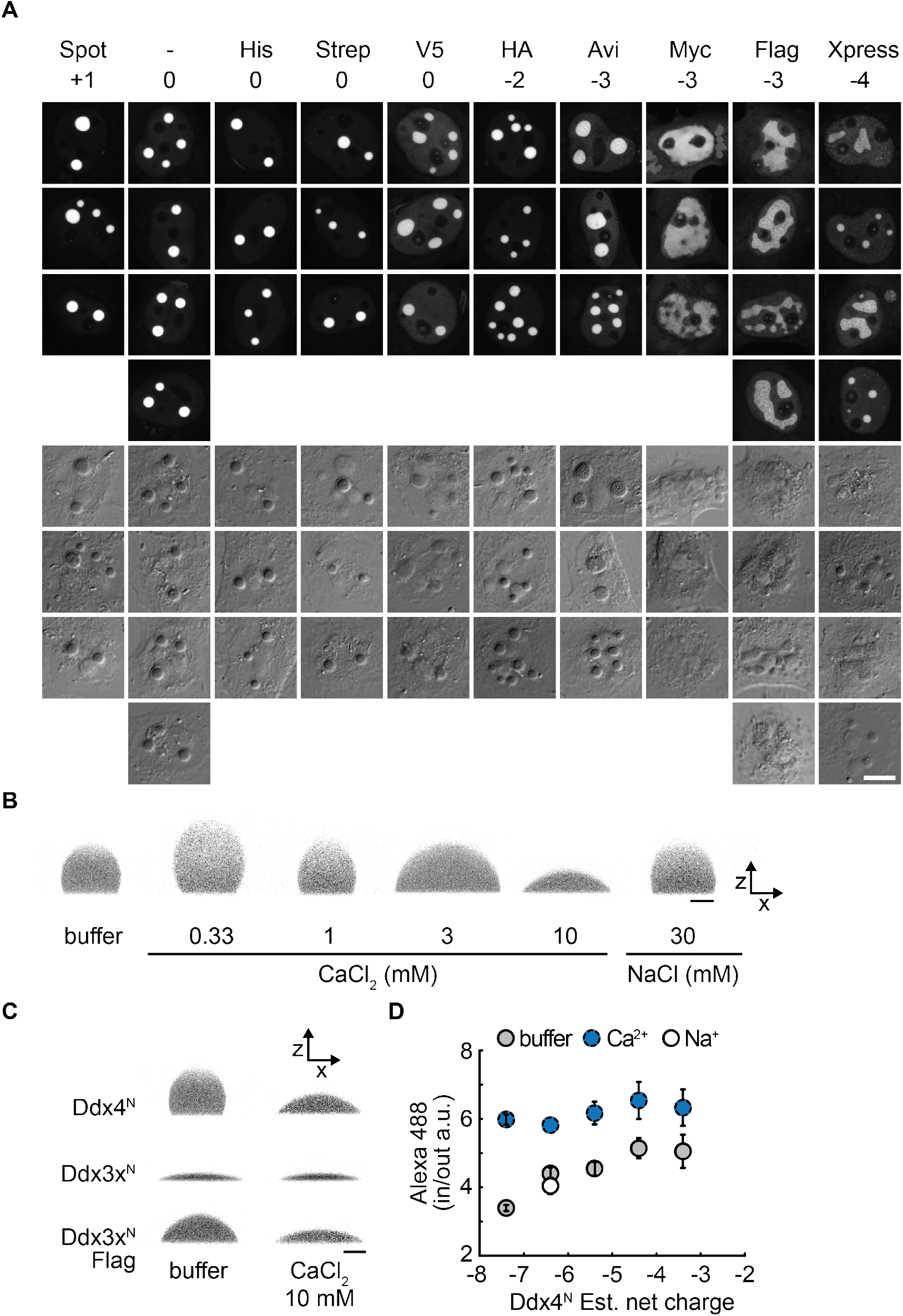
Net charge modulates the morphology and solvent properties of condensates. **(A)** Fluorescent (top) and DIC images (bottom) of HeLa cells expressing Ddx4^YFP^(−11.4) constructs. Addition of negatively charged tags to the C-terminus of the construct resulted in changes in condensate morphology, with the dense phase appearing less spherical. Analysis of the fluorescence intensity (Fig S1D) also indicated that Ddx4^YFP^ intensity in both the nucleoplasm and condensates varied upon addition of various C-terminal tags (tag sequences shown in Fig S1C). **(B)** Addition of CaCl_2_ resulted in titratable changes in the morphology of Ddx4^N^ condensates on siliconised glass coverslips, whereas addition of an equivalent ionic strength of NaCl did not. **(C)** Condensates composed of the less charged Ddx3x^N^ displayed a more wetted morphology compared to its Ddx4^N^ paralogue (estimated net charge of −6.4) and the morphology was unaltered after addition of 10 mM CaCl_2_. A more beaded profile and sensitivity to calcium was engineered into the Ddx3x^N^ construct through addition of a Flag tag that increased the estimated net charge from −1.1 to −4.1, indicating that the sensitivity to calcium was modulated by the excess charge. **(D)** The fluorescence intensity ratio of alexa 488 inside:outside of Ddx4^N^ condensates varied with the net charge of the construct. Addition of 10 mM CaCl_2_ increased the fluorescence ratio, with all constructs displaying similar values in the presence of 10 mM CaCl_2_. Addition of an equivalent ionic strength of NaCl indicated that this was not due to an ionic strength effect. Error bars represent the standard deviation.

**Figure S4.**
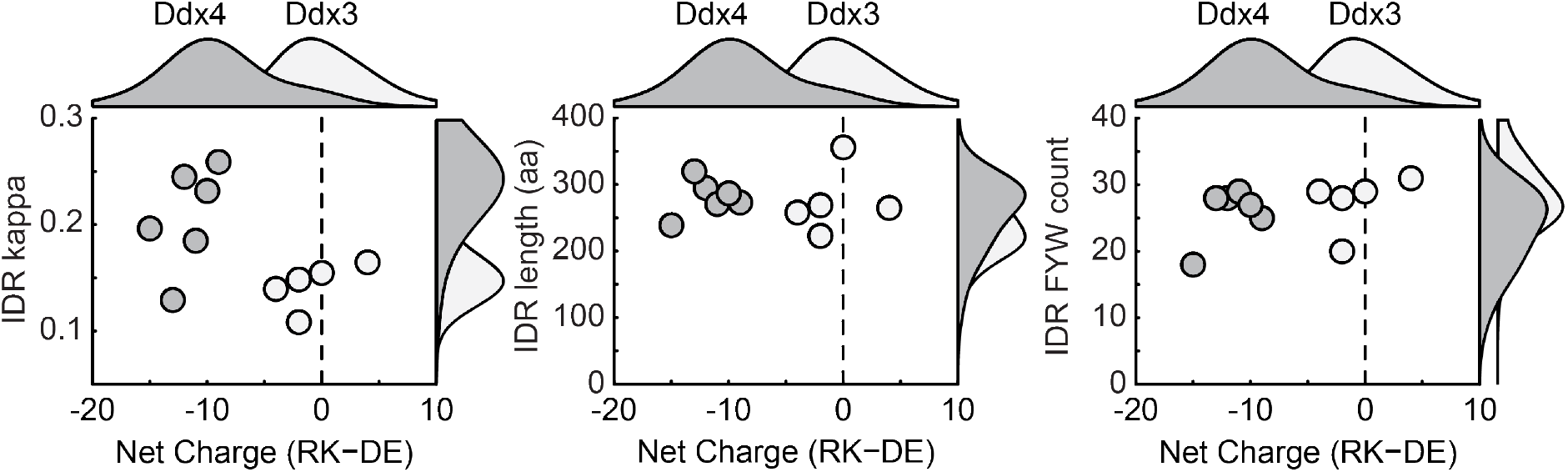
Ddx4 and its somatic paralogue, Ddx3x, differ in net charge and charge patterning. Density distributions represent data from Ddx4 (dark grey) and Ddx3x (light grey) orthologues annotated on Ensembl and indicate a difference in full-length net charge, and charge patterning in intrinsically disordered regions (IDR) between Ddx4 and Ddx3x sequences. In contrast, both IDR length and FYW (Phe, Tyr, Trp) count are similar for the two paralogues. Data for a comparatively phylogenetically broad subset of human, mouse, frog, fish, fruit fly and nemotode sequences are shown as individual points, and display the same trend. Previously, we showed that the charge patterning of Ddx4^N^ promoted its phase separation [24]. The conservation of repulsive long-range electrostatic interactions in Ddx4 therefore appears to be compensated by modulating an additional sequence feature that promotes phase separation.

